# HBs-S antigen-dependent enhancement of HBV infection

**DOI:** 10.1101/2022.06.07.495222

**Authors:** Takahiro Sanada, Osamu Yoshida, Yoichi Hiasa, Michinori Kohara

**Author notes:** Correspondence: Michinori Kohara, Department of Microbiology and Cell Biology, Tokyo Metropolitan Institute of Medical Science, 2-1-6, Kamikitazawa, Setagaya-ku, Tokyo 156-8506, Japan.

## Abstract

**Background Aims:** In natural infections with hepatitis B virus (HBV), large amounts of hepatitis B surface-small antigen (HBs-S) subviral particles (SVPs), which do not contain viral nucleocapsid, are secreted into the blood. The function of excess amounts of SVPs remains largely unknown. In this study, we analyzed the function of HBs-S in HBV infection.

**Methods:** The effect of HBs-S in HBV infection was evaluated in a human hepatoma cell line, in primary human hepatocytes, and in chimeric mice with humanized livers. To analyze the involvement of glycosaminoglycan, HBs-S attachment to cells in the presence of heparin was evaluated by enzyme-linked immunosorbent assay (ELISA), and the enhancement of viral attachment was evaluated by measurement of viral DNA. Additionally, the interaction between HBs-S and viral particles was analyzed by immunoprecipitation.

**Results:** Attachment of the HBV DNA to cells inoculated with the combination of virus and HBs-S was significantly higher than that to cells inoculated with HBV only. Pretreatment of the cells with HBs-S also increased viral DNA levels significantly compared to that in untreated cells. In contrast, HBs-L did not enhance the viral attachment. Enhancement of viral attachment was associated with the attachment of HBs-S to the cells via heparan sulfate. HBs-S also interacted with the viral particle. Furthermore, in chimeric mice with humanized livers, HBs-S enhanced HBV infection.

**Conclusions:** We demonstrated that HBs-S enhances viral attachment *in vitro* and viral infection *in vivo*. HBs-S interacted with heparan sulfate on the cellular surface, and this interaction contributed to the enhancement of viral attachment. These data indicate that HBs-S enhances the viral infection, and may contribute to the high transmissibility of HBV.

## Introduction

Infection by hepatitis B virus (HBV) causes acute and chronic hepatitis, and chronic infection can lead to cirrhosis and hepatocellular carcinoma. According to a report of the World Health Organization, 296 million people were living with chronic HBV infection in 2019, with 1.5 million new infections and 820 thousand people succumbing to HBV-related diseases each year [1]. Although an effective vaccine is available, HBV infection remains a major public health concern worldwide [2].

The main route of HBV infection is mother-to-child transmission, but, especially in adulthood, HBV infection also occurs by contact with infected blood and body fluids, such as needlestick injuries and sexual contact. Prospective studies of health care workers have estimated that the average risk of HBV transmission after a needlestick exposure is 30%, a risk higher than that of other bloodborne pathogen such as human immunodeficiency virus (0.3%) and hepatitis C virus (1.8%) [3]. It remains unclear why HBV has such high transmissibility.

The mature infectious HBV viral particle, called the “Dane particle”, is 42 nm in diameter and consists of an envelope and nucleocapsid containing HBV DNA and polymerase. The viral envelope consists of three different proteins, designated hepatitis B surface (HBs)-small (S),-medium (M), and -large (L). The S domain is common to all three proteins, whereas HBs-M has a pre-S2 domain, and HBs-L has both pre-S1 and pre-S2 domains. Since the pre-S1 domain mediates the viral interaction with the host cellular receptor, the sodium taurocholate cotransporting polypeptide (NTCP), HBs-L is the main component of viral particle [4].

In natural infection with HBV, large amounts of HBs-S antigen subviral particles (SVPs), which do not contain viral nucleocapsid, are secreted into the blood. SVPs have two main forms: 25-nm-diameter spheres, and 22-nm-diameter tubular filaments of variable length. The concentration of SVPs in HBV patient serum is 1,000 to 100,000 times higher than that of infectious particles [5]. The function of such excess amounts of SVPs remains largely unknown. Rydell *et al*. have shown that SVPs act as decoys to allow viral particles to escape neutralization by host antibodies [6]. In the duck hepatitis B virus (DHBV) model, SVPs have been reported to both enhance and inhibit cell attachment by the virus [7, 8].

In the present study, we focused on the function of HBs in HBV infection. Using a human hepatoma cell line, primary human hepatocytes, and chimeric mice with humanized livers, we analyzed the role of HBs in HBV infection.

## Materials and Methods

### Viruses

Inocula of HBV genotype C (C_JPNAT; Accession No.: AB246345.1) were obtained from the supernatant of primary human hepatocytes derived from chimeric mice with humanized livers (PXB-cells; PhoenixBio, Hiroshima, Japan) at 8-10 weeks after infection.

### Human sera

Sera from four donors were used in this study (Table 1). Two donors were chronic hepatitis B patients treated with nucleos(t)ide analogues (NAs). The other two donors were chronic asymptomatic HBV carriers (ASCs) who were not undergoing anti-HBV treatment. Three of the donors are infected with genotype-C HBV; the fourth is infected with genotype-D virus. All donors provided written informed consent. The clinical trial was approved by the Certified Review Board in Ehime University (ref 18EC003), registered in UMIN Clinical Trials Registry (ref UMIN000027442) and the Japan Registry of Clinical Trials (ref jRCTs061180100), and conducted in accordance with the principles of the Declaration of Helsinki and Good Clinical Practice. AB human serum (Pel-Freez Biologicals, Rogers, AR, USA) was used as control serum.

**Table 1.** Characterization of HBV patient sera used in this study.

| Patient ID | sex | age | HBV genotype | HBsAg (IU/mL) | HBV-DNA (Log IU/mL) | HBeAg (COI) | HBsAb | HBeAb | Remarks |
| --- | --- | --- | --- | --- | --- | --- | --- | --- | --- |
| NAs #1 | Male | 59 | C | 4213 | 0 | 0 | Neg | Pos | Under treatment with entecavir |
| NAs #2 | Female | 72 | C | 0.18 | 0 | 0 | Neg | Neg | Under treatment with entecavir |
| ASC #1 | Female | 43 | D | 1527 | 2 | 0 | Neg | Pos | - |
| ASC #2 | Male | 65 | C | 48 | <1.3 | 0 | Neg | Pos | - |

COI: cut off index
Pos: positive, Neg: negative

### Animals

Chimeric mice with humanized livers (PXB-mouse) were purchased from PhoenixBio.

### Cells

HepG2 and HepG2-hNTCP-30 cells were cultured on type I, collagen-coated, 48-well plates as described previously [9]. PXB-cells were cultured on type I, collagen-coated, 48-well plates using maintenance medium as previously described [10].

### Viral protein

Purified HBs-S, and HBs-L proteins were purchased from Beacle (Kyoto, Japan). These proteins were obtained with sequences corresponding to those of the genotype-C virus. Purified HBs-S and HBs-L proteins were provided as proteins expressed in yeast. According to the manufacturer, the viral proteins form particle-like structures. HBs protein derived from human serum also was purchased from Beacle. The HBs antigen subtype was ay.

Biotinylation of HBs-S and HBs-L was performed using the Antibody Biotinylation Kit for IP (Thermo Fisher Scientific, Waltham, MA, USA) according to the manufacturer’s instructions.

### Serological test of patient sera

The serological test for HBV was performed using a chemiluminescent immunoassay (Architect i2000SR; Abbott, Abbott Park, IL, USA), specifically targeting HBs antigen (HBsAg), hepatitis B e antigen (HBeAg), anti-HBs antibody (HBsAb), and anti-HBe antibody (HBeAb). Measurement of viral DNA was performed using a quantitative polymerase chain reaction (qPCR) assay (Cobas 6800; Roche, Basel, Switzerland).

### Ultracentrifugation

To remove infectious viral particles from serum, the sera of HBV patients were centrifuged at 96,000 × g for 5 h at 4°C as described previously [11]. The resulting supernatants were used for experiments.

### Viral infection of chimeric mice with humanized livers

All animal experiments were performed by PhoenixBio. HBV (2.0 × 10^5^ viral DNA copies/mL) was mixed with an equal volume of each ultracentrifuged serum sample; the mixtures then were incubated for 1 h at room temperature. Aliquots of the resulting suspensions were inoculated into chimeric mice with humanized livers at 100 μL (1.0 × 10^4^ viral DNA copies) per animal. Serum samples then were collected from the infected mice each week. Viral DNA from the serum samples was analyzed by qPCR as described previously [12].

For analysis of the effect of HBs-S, HBV (2.0 × 10^5^ viral DNA copies/mL) was mixed with an equal volume of control serum sample containing HBs-S (20 μg/mL); the mixture then were incubated for 1 h at room temperature. Aliquots of the resulting suspensions were inoculated into chimeric mice with humanized livers at 100 μL (1.0 × 10^4^ viral DNA copies) per animal. Serum samples then were collected from the infected mice each week and used for analysis.

### Attachment assay

HBV inocula containing 3.75 × 10^5^ viral DNA copies with varying amounts (0.1, 1, or 10 μg/mL) of viral protein were incubated at 37°C for 1 h. An aliquot (125 μL) of each mixture then was used to inoculate an aliquot (containing 5.0 × 10^4^ cells) of the HepG2-hNTCP-30 cell line or the PXB-cells yielding a multiplicity of infection of 7.5 genome equivalent (GEq)/cell. After incubation for 3 h on ice, the cells were washed 5 times with culture medium and then collected at 400 × g for 5 min at 4°C. The resulting cell pellets were used for qPCR analysis as described previously [13]. The viral DNA of a sample infected with HBV only was used as the control.

Pretreatment (1 h) with HBs-S or HBs-L (0.1, 1, or 10 μg/mL) or with biotin-conjugated HBs-S or HBs-L (1 μg/mL) was performed prior to viral infection. The cells were washed 5 times with culture medium and then inoculated with 7.5 GEq/cell of HBV. After incubation for 3 h on ice, the cells were washed 5 times with culture medium and then collected at 400 × g for 5 min at 4°C. The resulting cell pellets were used for qPCR analysis as described previously [13]. The viral DNA of a sample infected with HBV (no HBs pretreatment) was used as the control.

For analysis of the effect of heparin, HBs-S or HBs-L diluted to 10 μg/mL was mixed with the heparin (0-1,000 μg/mL) and incubated at 37°C for 1 h. Aliquots (125 μL/well) of each mixture then were used to inoculate the wells of a plate containing HepG2-hNTCP-30 cells (5.0 × 10^4^ cells/well) and the plates were incubated at 37°C for 1 h. After washing 5 times with culture medium, the cells were infected with HBV and then collected for analysis as described above.

### Enzyme-linked immunosorbent assay (ELISA) of HBs-S attachment

HepG2-hNTCP-30 cells (2.0 × 10^4^ cells/well) were cultured in type I collagen-coated, 96-well plates. After overnight incubation at 37°C, the cells were fixed with 4% paraformaldehyde for 20 min at room temperature. After washing with phosphate-buffered saline (PBS), fixed cells were blocked by addition of blocking buffer (PBS containing 1% bovine serum albumin, 0.5% Tween 20, and 2.5 mM ethylenediaminetetraacetic acid, EDTA) at 200 μL/well followed by incubation at room temperature for 2 h. Next, a solution of blocking buffer containing HBs-S diluted to 1 μg/mL and heparin diluted to 0-1,000 μg/mL was distributed to the plates at 50 μL/well, and the plates were incubated at room temperature for 1 h. After the plates were washed three times with PBS containing 0.05% Tween 20 (PBST), a solution of 50 μL of 1 μg/mL anti-HBs-S mouse monoclonal antibody (81A22-6; raised in-house) in blocking buffer was distributed to the wells, and the plates were incubated at room temperature for 1 h. After the plates were washed three times with PBST, peroxidase-conjugated anti-mouse-IgG antibody (GE Healthcare, Chicago, IL, USA), diluted 1:5,000 in blocking buffer, was distributed to the plates at 50 μL/well. After 1 h of incubation at room temperature, the plates were washed five times with PBST. Then, o-phenylenediamine substrate in hydrogen peroxide was distributed to the plates at 100 μL/well, and the plates were incubated at room temperature for 20 min. The reactions were stopped by the addition of 50 μL/well of 2 M sulfuric acid and absorbance was measured at 492 nm.

### Immunoprecipitation

Spent culture medium (200 μL at 2.0 × 10^6^ copies/mL) was mixed with an equal volume of Dulbecco’s Modified Eagle Medium containing 2 μg/mL biotin-conjugated HBs-S or HBs-L. The mixture was incubated at 37°C for 1 h. Each mixture then was combined with an equal volume of binding buffer (20 mM Tris-HCl (pH 7.4), 1 mM EDTA, 2 M NaCl, 0.1% Tween 20). The resulting mixture was combined with 100 μL of streptavidin-conjugated magnetic beads (MS160/Streptavidin; JSR Life Sciences, Ibaraki, Japan), and the combined volume was incubated with rotation at room temperature for 10 min. The mixture then was vortexed and placed on a magnetic stand for 1 min, the supernatant was decanted, and the pellets were washed three times with binding buffer. The resulting pellets were used for qPCR analysis as described previously [13].

### Statistical analyses

Statistical analyses were performed with R software version 4.0.3 (https://www.r-project.org/). Data were analyzed using two-tailed Welch’s t-test and Dunnett’s test were used to conduct statistical analyses of the data. *p* values of less than 0.05 were considered significant.

## Results

### HBV patient serum enhances viral propagation in chimeric mice

Since HBV has high transmissibility, we hypothesized that HBV patient serum contains a factor that enhances viral infection. To analyze the effect of HBV patient serum on viral infection, we inoculated chimeric mouse with humanized livers with a mixture of HBV and HBV patient serum (Fig. 1A). To remove infectious viral particles from HBV patient serum samples, the sera were ultracentrifuged and the resulting supernatants were used for this experiment. All supernatant samples were confirmed as HBV-DNA negative. When HBV was mixed with the samples from HBV patients who received NAs or from ASCs, the viral DNA was detected in the sera of the inoculated chimeric mice earlier than in control samples (Fig. 1B). This result indicated that HBV patient serum contains a factor that enhances viral infection.

**Figure 1.**
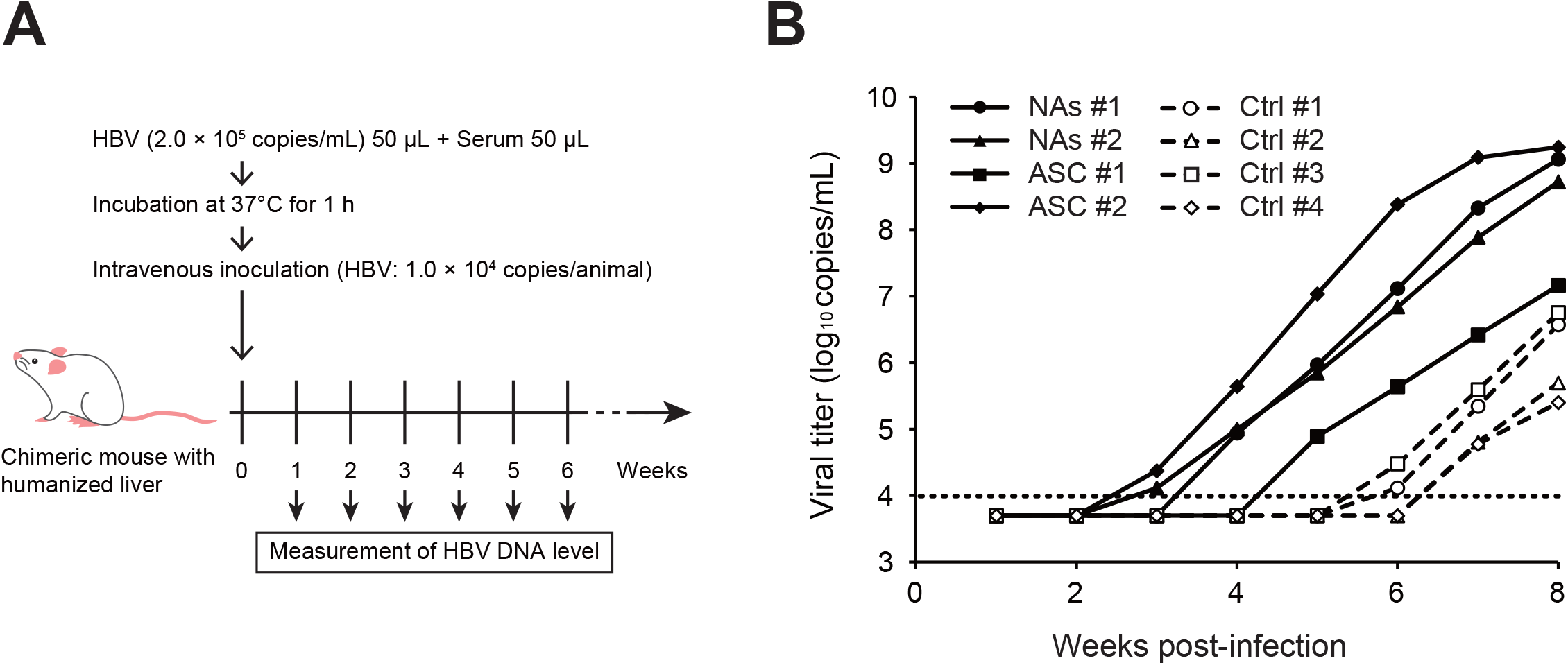
HBV propagation in chimeric mice with humanized livers following inoculation with mixtures of HBV and HBV patient serum. (A) Experimental schedule of HBV infection in chimeric mice with humanized livers. (B) Viral titer in serum samples collected from chimeric mice with humanized livers following inoculation with mixtures of HBV and HBV patient serum. Horizontal dotted line indicates the lower limit of detection (1.0 × 10^4^ copies/mL). Viral titers below the detection limit are plotted as 5.0 × 10^3^ copies/mL. NAs, chronic hepatitis B patients treated with nucleos(t)ide analogues; ASCs, chronic asymptomatic HBV carriers; Ctrl, control.

### HBs-S enhances HBV attachment to NTCP-expressing HepG2 cells

Since HBV patient serum contains large quantities of HBs-S antigen [5], we focused on this molecule as a possible mediator of this enhancement. We evaluated the effects of HBs-S and HBs-L in HBV attachment to NTCP-expressing HepG2 cells (HepG2-hNTCP-30 cells) [9]. The HBs-S or HBs-L protein was mixed with HBV, and the resulting mixture then was inoculated to the cells (Fig. 2A). After 3 h of incubation on ice, Dane particle attachment to the cells was evaluated by measurement of viral DNA. Attachment of the HBV DNA to cells inoculated with a mixture of HBV and HBs-S was significantly higher than that to cells inoculated with HBV only (Fig. 2B). In contrast, there was no significant difference in attachment between cells inoculated with a mixture of HBV and HBs-L and cells inoculated with HBV only. Notably, pretreatment of the cells with HBs-S also significantly increased viral DNA levels compared to cells that were not pretreated, while pretreatment with HBs-L did not provide this enhancement (Fig. 2A and C). These data indicated that HBs-S, and not HBs-L, enhances viral attachment to NTCP-expressing cells.

**Figure 2.**
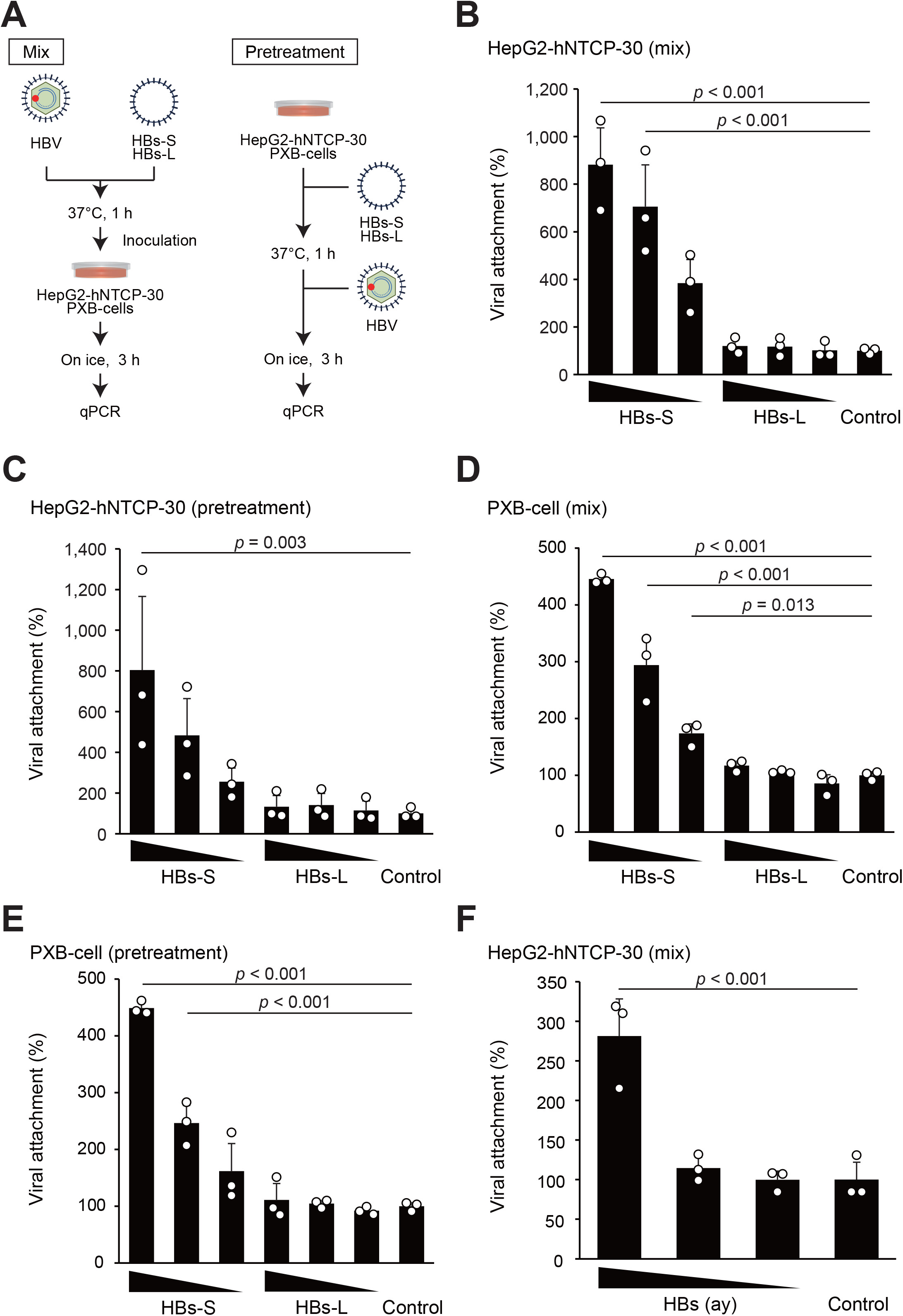
Enhancement of HBV attachment *in vitro*. (A) Schematic diagram of HBV attachment experiment. (B) HBV DNA levels of HepG2-hNTCP-30 cells following inoculation with HBV in combination with either protein. (C) HBV DNA levels of HepG2-hNTCP-30 cells following pretreatment with either protein prior to inoculation with HBV. (D) HBV DNA levels of PXB-cells inoculated with HBV in combination with either protein. (E) HBV DNA levels of PXB-cells following pretreatment with either protein prior to inoculation with HBV. (F) HBV DNA levels of HepG2-hNTCP-30 cells following inoculation with HBV and HBs protein derived from HBV patients. Values are shown as mean ± SD of data from 3 independent experiments per group. *p* values were calculated using a two-tailed Dunnett’s test. Where not indicated, differences were not significant (*p* ≥ 0.05).

Next, we repeated these experiments using primary human hepatocytes (PXB-cells). As seen with HepG2-hNTCP-30 cells, viral attachment to PXB-cells was significantly enhanced with the mixture of HBV and HBs-S, compared to that observed with HBV only (Fig. 2D). Additionally, pretreatment of PXB-cells with HBs-S cells significantly enhanced viral attachment, compared to that seen with HBV only (Fig. 2E). Thus, enhancement viral attachment by HBs-S also was observed in primary human hepatocytes.

The above assays used HBs-S and HBs-L that had been expressed in yeasts; to validate our results, we repeated the attachment experiments using HBs antigen proteins derived from HBV patients. This antigen seems to contain not only HBs-S, but also HBs-L. Attachment of the HBV DNA to cells inoculated with a mixture of HBV and HBs derived from HBV patients was significantly higher than that to cells inoculated with HBV only (Fig. 2F).

Considered together, these data indicated that HBs-S enhances the attachment of HBV to cells.

### Enhancement of HBV attachment by HBs-S is mediated by interaction with heparan sulfate

Pretreatment of cells with HBs-S enhanced viral attachment, suggesting that HBs-S interacts with the cellular surface. To address this hypothesis, we sought to identify the factor(s) that interact with HBs-S. To evaluate the involvement of the NTCP, the host cell receptor for HBV infection, we evaluated the viral attachment enhancement in HepG2 cells. Attachment analysis revealed that enhancement of viral attachment by HBs-S also was observed in HepG2 cells, as with HepG2-hNTCP-30 cells (Fig. 3A and B). Thus, some molecule(s) other than NTCP appear to contribute to the enhancement of viral attachment by HBs-S antigen.

**Figure 3.**
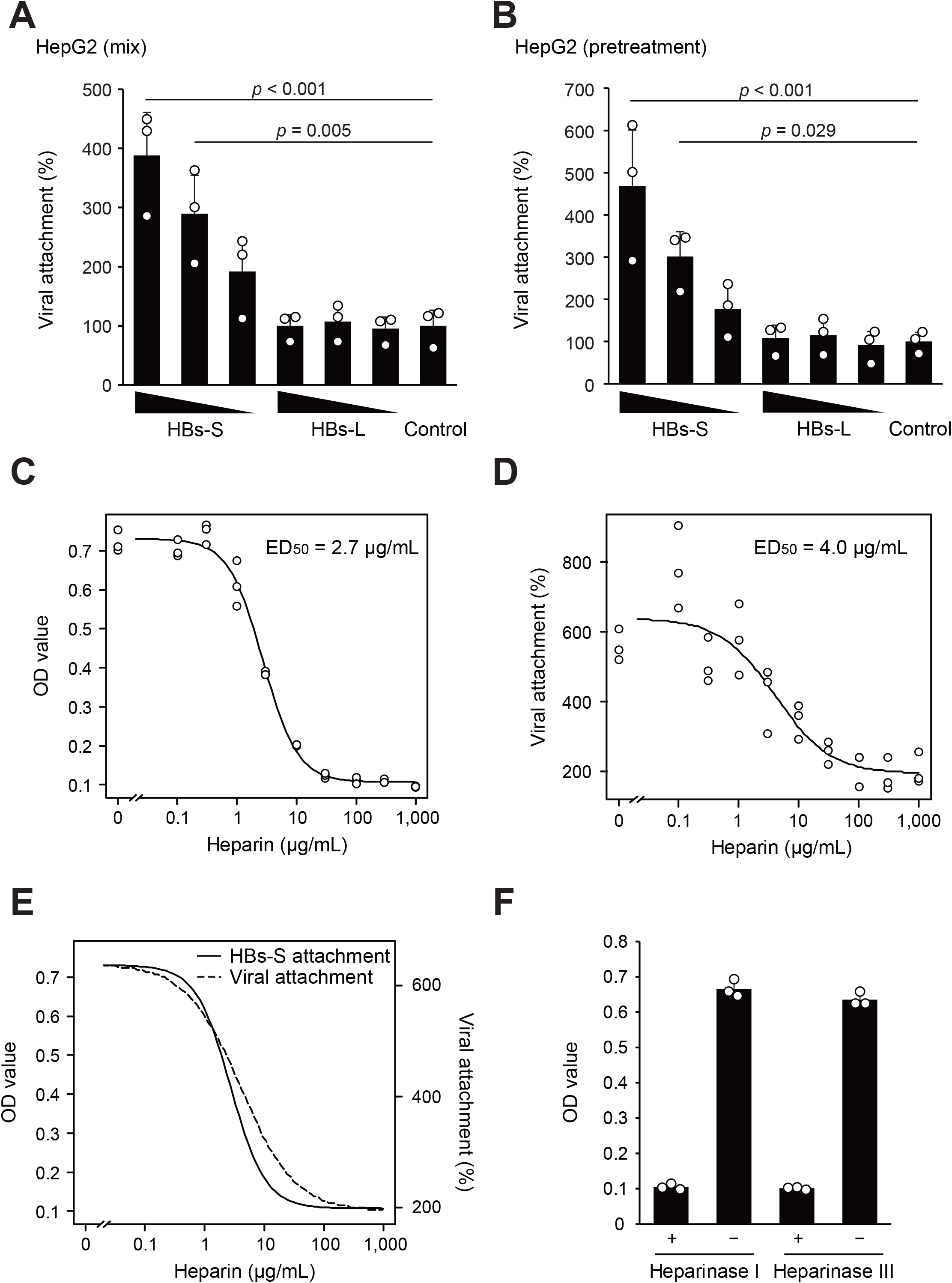
Involvement of heparan sulfate in enhancement of HBV attachment by HBs-S. (A) HBV DNA levels of HepG2 cells following inoculation with HBV in combination with either protein. (B) HBV DNA levels of HepG2 cells following pretreatment with either protein prior to inoculation with HBV. (C) HBs-S attachment to HepG2-hNTCP-30 cells at a range of concentrations of heparin, as assessed by ELISA. (D) HBV DNA levels of HepG2-hNTCP-30 cells pretreated with HBs-S and heparin prior to inoculation with HBV. (E) Dose-HBs-S attachment curve and dose-viral attachment curve. (F) Effects of heparinase I and III on HBs-S attachment to HepG2-hNTCP-30 cells. Values are shown as mean ± SD of data from 3 independent experiments per group. *p* values were calculated by a two-tailed Dunnett’s test. Where not indicated, differences were not significant (*p* ≥ 0.05).

The HBs-S protein carried by the Dane particle has been reported to interact with heparan sulfate proteoglycan [14]. Therefore, we analyzed whether HBs-S alone interacts with heparan sulfate and whether the HBs-S-mediated enhancement of viral attachment is associated with the interaction between HBs-S and heparan sulfate. First, we used ELISA to evaluate HBs-S attachment to cells. This assay revealed that HBs-S does indeed attach to cells, while premixing of HBs-S with heparin resulted in dose-dependent inhibition of the attachment HBs-S to cells (Fig. 3C). Next, we analyzed the effect of heparin on the enhancement of viral attachment by HBs-S, showing that heparin also inhibited HBs-S-mediated enhancement of viral attachment (Fig. 3D). Thus, HBs-S attachment to the cells correlated with the enhancement of viral attachment (Fig. 3E).

These data indicated that HBs-S attaches to cells via heparin and/or a heparin-like molecule. Next, we determined the cellular factor involved in HBs-S attachment. Heparinase I digests both heparin and heparan sulfate, and heparinase III specifically digests heparan sulfate. Both heparinase I and III treatments strongly inhibited the HBs-S attachment to the cells (Fig. 3F). Thus, HBs-S appears to attach to cells via heparan sulfate, as is known for the infectious viral particle.

### HBs-S interacts with the infectious viral particle

Next, we evaluated the interaction between HBs-S and the infectious viral particle. This experiment employed biotin-conjugated HBs-S and HBs-L. We confirmed that pretreatment of the cells with biotin-conjugated HBs-S increased viral attachment significantly compared with non-treatment (Fig. 4A). We then immunoprecipitated the infectious viral particle with biotin-conjugated HBs-S and HBs-L (Fig. 4B). The level of infectious viral particle that interacted with HBs-S or HBs-L was determined by measurement of viral DNA in the immunoprecipitated sample. The viral DNA level of specimens recovered from immunoprecipitation with HBs-S was significantly higher than that of control specimens (Fig. 4C). In contrast, the viral DNA level of specimens recovered from immunoprecipitation with HBs-L did not differ significantly from that of control specimens. These results indicated that HBs-S interacts with infectious viral particles.

**Figure 4.**
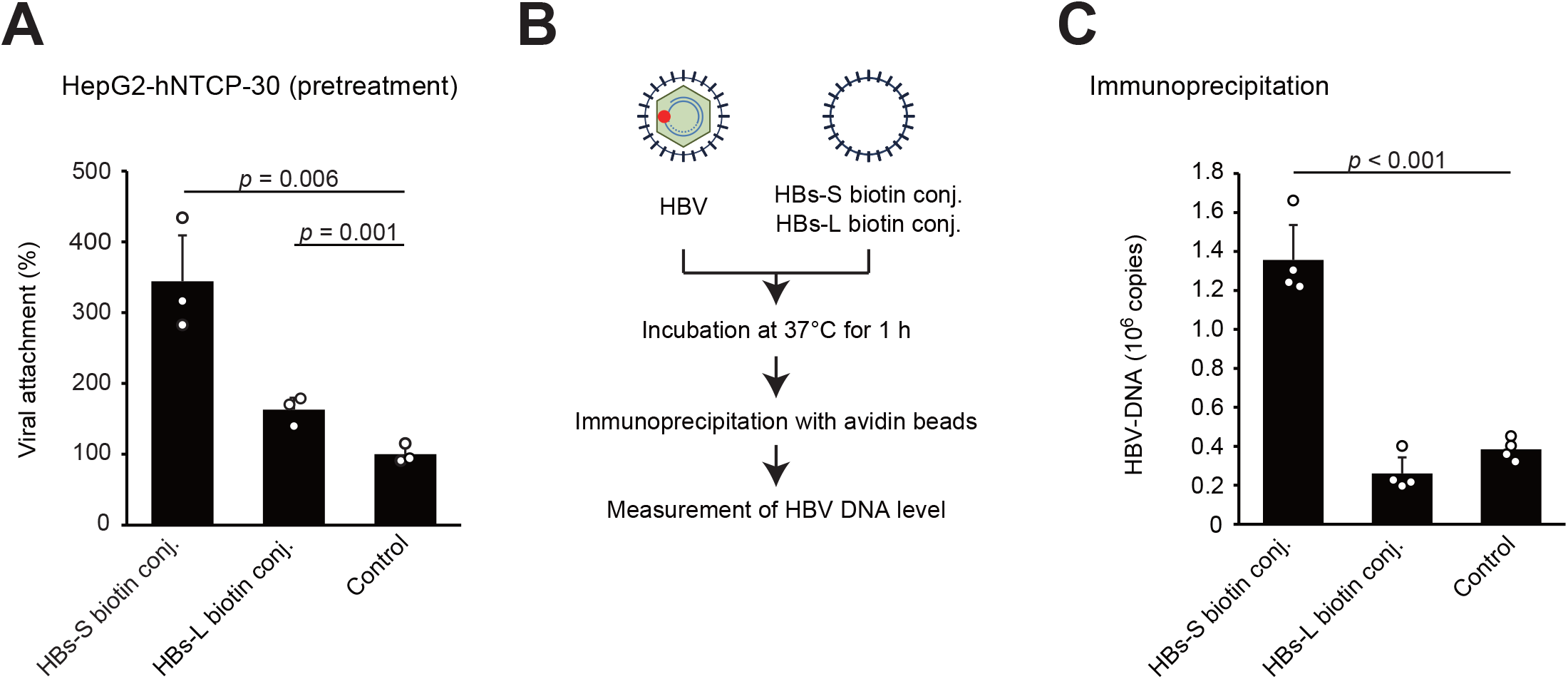
Interaction between HBs-S and infectious viral particles. (A) HBV DNA levels of HepG2-hNTCP-30 cells treated with either protein prior to inoculation with HBV. (B) Schematic diagram of immunoprecipitation experiment. (C) HBV DNA levels after immunoprecipitation of HBV mixed with either protein. Values are shown as mean ± SD of data from 3 or 4 independent experiments per group. *p* values were calculated by a two-tailed Dunnett’s test. Where not indicated, differences were not significant (*p* ≥ 0.05).

### HBs-S enhances HBV infection in chimeric mice with humanized livers

Finally, we evaluated the effect of HBs-S on HBV infection in chimeric mice with humanized livers. HBV with or without HBs-S was used to inoculate the chimeric mice, and the serum viral DNA level then was evaluated (Fig. 5A). In chimeric mice infected with the combination of HBV with HBs-S, viral DNA was detected starting from 4 weeks post-infection; compared to the viral DNA levels in chimeric mice infected with HBV only, the viral DNA levels in chimeric mice infected with HBV with HBs-S were significantly higher at 5 to 7 weeks post-infection (Fig. 5B). These data indicated that HBs-S enhances HBV infection *in vivo*.

**Figure 5.**
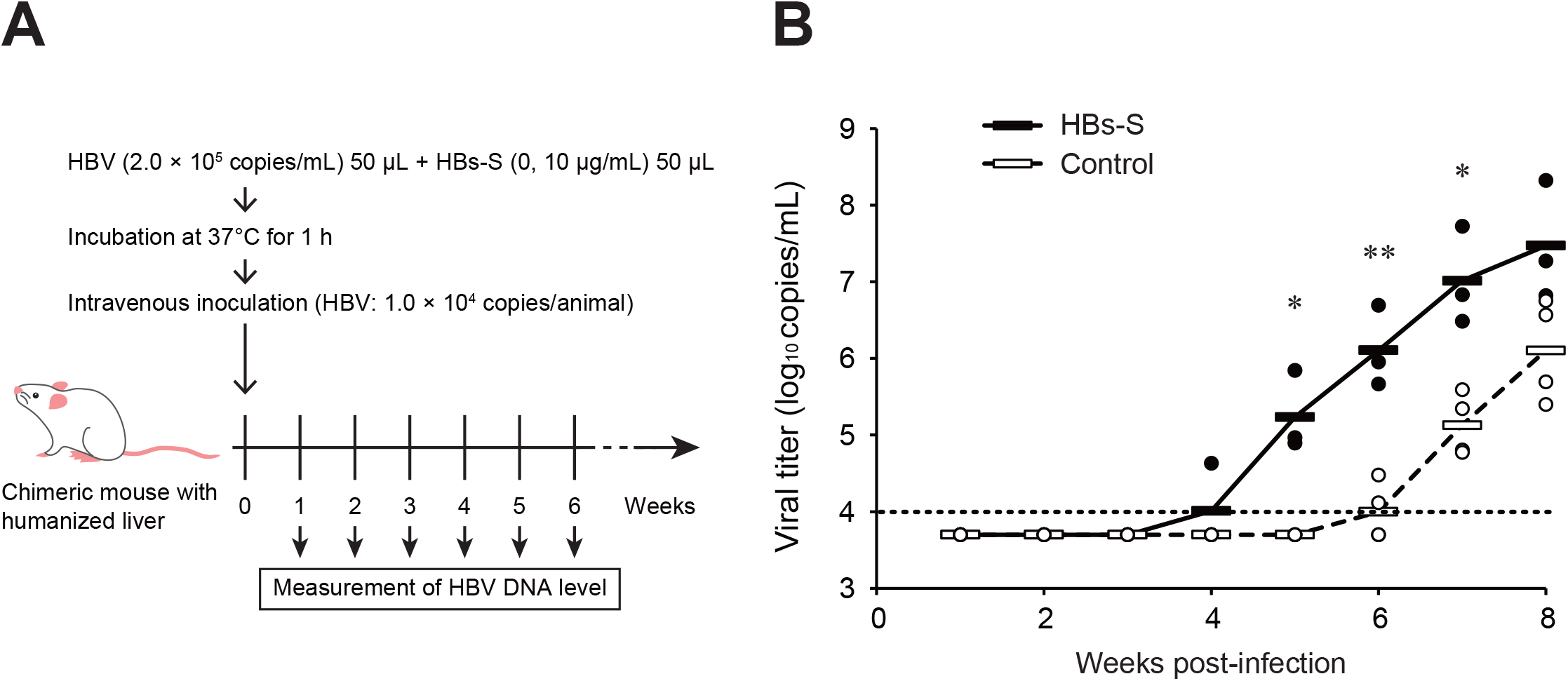
HBV propagation in chimeric mice with humanized livers following inoculation with a mixture of HBV and HBs-S. (A) Experimental schedule of HBV infection in chimeric mice with humanized livers. (B) Viral titer in serum samples collected from chimeric mice with humanized livers following infection with HBV with or without HBs-S. Horizontal dotted line indicates the lower limit of detection (1.0 × 10^4^ copies/mL). Viral titers below the detection limit are plotted as 5.0 × 10^3^ copies/mL. Thick horizontal bars indicate arithmetic mean values in each group for data from 3 animals per group. Significant differences were calculated by a two-tailed Welch’s t-test (\**p* < 0.05, \*\**p* < 0.01). Where not indicated, differences were not significant (*p* ≥ 0.05).

## Discussion

Large amounts of HBs-S are secreted into the blood in HBV patients, but the role of this secreted protein remains still largely unknown [5]. One of the known functions of HBs-S is to serve as a decoy for host-generated neutralizing antibodies [6]. Specifically, SVPs of HBs-S interact with neutralizing antibody, thereby protecting Dane particles from the humoral immune response. This approach is an effective strategy to facilitate infection in people who already possess an immune response against HBV or who are sustaining a chronic infection by induction of immune tolerance. However, this observation is not sufficient to explain the high transmissibility of HBV. The present study showed that HBs-S enhances the viral infection in a dose-dependent manner. This analysis employed a maximum HBs-S concentration of 10 μg/mL. According to the manufacturer, a concentration of 10 μg/mL HBs-S is equivalent to 5,000-10,000 IU/mL. For comparison, the HBs antigen (HBsAg) level in HBV patients can be exceed 100,000 IU/mL [15]; thus, the enhancement of HBV attachment in patient serum may exceed that detected in the experiments described here. In actuality, infection by HBV is the result of contact with infected blood and body fluids, such as needlestick injury and sexual contact. HBV can be transmitted when a small amount of infected blood or body fluid enters the circulatory system of an HBV-naïve recipient; under such conditions, the infecting blood or body fluids will be diluted, and the concentration of donor’s HBs-S in the recipient’s blood will become extremely low. In our experiment, chimeric mice with humanized livers were inoculated with a mixture of HBV and 10 μg/mL HBs-S; conditions under which the HBs-S entering the bloodstream would be greatly diluted. Nonetheless, we observed that HBs-S enhanced HBV infection in this chimeric mouse model. Our study also showed that HBs-S interacts with both heparan sulfate and the Dane particle. Thus, the interaction of HBV with HBs-S may facilitate attachment of the HBV particle to hepatocytes, such that HBs-S acts as a bridge between the Dane particle and heparan sulfate or other cellular receptors such as NTCP.

The present study showed that HBs-S interacts with heparan sulfate, a component of the cellular surface. The Dane particle also has been shown to interact with heparan sulfate, thereby attaching to hepatocytes [16]. Notably, however, HBs-S did not appear to interfere in this attachment process, as might be expected if the Dane particle and HBs-S compete for the same binding site(s) on heparan sulfate. A previous study also reported that SVPs do not inhibit the ability of HBV to infect primary human hepatocytes [17]. We postulate that the Dane particle may attach either to heparan sulfate on the cellular surface or to HBs-S attached to the cells, subsequently moving to the NTCP receptor.

Previously, Bruns *et al*. reported that, in DHBV, SVP enhanced DHBV infection [7]. Those authors showed that SVP enhances intracellular viral replication and gene expression. Bruns *et al*. focused on post-infection stages and speculated that the induction of intracellular signaling events was occurring. Based on those results, we infer that HBs-S does not function solely to enhance viral attachment to cells, but likely plays multiple roles in the course of HBV infection. Additionally, since HBV patient serum enhanced HBV infection in chimeric mice with humanized livers (as shown in the present study), we expect that viral protein other than HBs-S (e.g., HBeAg or other serum components) also contribute to the enhancement of HBV infection. To test these hypotheses, we are undertaking further analyses to clarify the mechanism of the high transmissibility of HBV

In the present study, both HBsAg-high patient serum and HBsAg-low patient serum (0.18 IU/mL) provided enhancement of HBV infection (Fig. 1B). Previously, the literature has reported the existence of immunocomplexes of HBsAbs and HBsAgs in the circulating blood of HBV patients [18, 19]. Since clinical assays for HBsAg detection generally are based on immunoreaction between antibodies and HBsAgs, the actual HBsAg levels of HBV patients may be higher than the measured values. Using the dissociation of HBsAg from HBsAg-HBsAb immunocomplexes, Matsumoto *et al*. also reported the detection of HBsAg in samples that tested HBsAg-negative by conventional assay [19]. Considered together, these studies suggest that HBV patients may possess blood HBsAg at levels exceeding the measured content, even in nominally HBsAg-negative HBV cases. HBV infection enhancement by HBs-S may be occurring in the sera of many or all HBV patient, and this phenomenon may facilitate the maintenance of HBV in individual patients as well as the spread of HBV among patients.

In conclusion, we demonstrated that HBs-S enhances HBV viral attachment *in vitro* and viral infection *in vivo*. HBs-S was shown to interact with heparan sulfate on the cellular surface, and this interaction contributed to the enhancement of viral attachment. The ability of HBs-S to enhance viral infection may contribute to the high transmissibility of HBV.

## Abbreviations

ASC: chronic asymptomatic carriers
DHBV: duck hepatitis B virus
EDTA: ethylenediaminetetraacetic acid
ELISA: enzyme-linked immunosorbent assay
GEq: genome equivalent
HBeAb: anti-hepatitis B e antibody
HBeAg: hepatitis B e antigen
HBsAb: anti-hepatitis B surface antibody
HBsAg: hepatitis B surface antigen
HBs-S: hepatitis B surface small
HBs-M: hepatitis B surface medium
HBs-L: hepatitis B surface large
HBV: hepatitis B virus
NA: nucleos(t)ide analogue
NTCP: sodium taurocholate cotransporting polypeptide
PBS: phosphate-buffered saline
PBST: phosphate-buffered saline containing 0.05% Tween20
qPCR: quantitative polymerase chain reaction
SVP: subviral particle

## Conflict of interest statements

The authors declare no conflict of interest.

## Financial support statement

This work was supported by JSPS KAKENHI Grant Number 19K16680 and AMED Grant Number JP20fk0210033.

## Authors’ contribution

Concept and design: T.S. and M.K. Acquisition of data: T.S. and O.Y. Analysis and interpretation of data: T.S. and M.K. Drafting of the manuscript: T.S. and M.K. Critical revision of the manuscript for important intellectual content: T.S., O.Y., Y.H. and M.K.

## Acknowledgements

We are grateful to all of the members of Department of Microbiology and Cell Biology. Tokyo Metropolitan Institute of Medical Science.

